# A Self-Healing, Viscoelastic Hydrogel Promotes Healing of Brain Lesions

**DOI:** 10.1101/2022.05.05.490746

**Authors:** Yan Hu, Yuanbo Jia, Siwei Wang, Yufei Ma, Guoyou Huang, Tan Ding, Dayun Feng, Guy M. Genin, Zhao Wei, Feng Xu

## Abstract

Brain lesions can arise from traumatic brain injury, infection, and craniotomy. Although injectable hydrogels show promise for promoting healing of lesions and health of surrounding tissue, enabling cellular ingrowth and restoring neural tissue continue to be challenging. We hypothesized that these challenges arise in part from viscoelastic mismatch between the hydrogel and the brain parenchyma, and tested this hypothesis by developing and evaluating a self-healing hydrogel that mimicked both the composition and viscoelasticity of native brain parenchyma. The hydrogel was crosslinked by dynamic boronate ester bonds between phenylboronic acid grafted hyaluronic acid (HA-PBA) and dopamine grafted gelatin (Gel-Dopa). This HA-PBA/Gel-Dopa hydrogel could be injected into a lesion cavity in a shear-thinning manner with rapid hemostasis, high tissue adhesion and efficient self-healing. We tested this in an *in vivo* mouse model of brain lesions and found the hydrogel to support neural cell infiltration, decrease astrogliosis and glial scars, and close the lesions. The results suggest a role for viscoelasticity in brain lesion healing, and motivate additional experimentation in larger animals as the technology progresses towards potential application in humans.

## 1. Introduction

Lesions and abscesses in the brain can arise following infection or traumatic brain injury (TBI) ^[1]^. The resulting cavities lack the extracellular matrix (ECM) scaffold needed to promote cellular infiltration, axonal extension, and neural regeneration required for healing ^[2]^, and instead promote glial scars through recruitment and activation of glial cells ^[3]^. Glial scars maintain homeostasis immediately following TBI, but prevent healing of the cavity in the long term through increased ECM deposition ^[2, 4]^ and secretion of inhibitory molecules ^[5]^. Efforts to develop material systems that inhibit glial scar formation and promote neural regeneration is thus a focus of a growing body of literature.

Previous efforts involve filling a lesion cavity with functional hydrogels that can provide mechanical support with a modulus in the range of 1-400 kPa ^[6]^. Although some studies in the literature purport to show that these materials heal brain injury, a critical look at this literature must note that no animal behavioral studies to support such claims have ever been performed, and that fundamental barriers persist. These barriers include developing technologies to avoid inducing astrogliosis, and fostering ingrowth and connectivity of axons. Overcoming these barriers is thus the focus of this work.

Our efforts were motivated by observations that many cells spread more when cultured in three-dimensional (3D) hydrogels that are viscoelastic than in hydrogels that are elastic ^[7]^, and that brain parenchyma is soft and viscoplastic ^[7-8]^. We sought to develop a self-healing hydrogel with the low stiffness and fast stress relaxation of brain parenchyma, which could be injected into a brain lesion with *in situ* solidification (sol-gel transitions) with controllable material dispersion within the brain.

Our approach centered on a hydrogel with a dopamine (Dopa) modified gelatin (Gel-Dopa) backbone crosslinked by phenylboronic acid (PBA) modified hyaluronic acid (HA-PBA). Gelatin and HA were selected because gelatin has bioactivity and cell adhesion sites, while HA is the major structural and bioactive component of native brain ECM that has been used previously in brain lesions ^[9]^. The dynamic boronate ester bonds arising from PBA/Dopa reactions result in tunable viscoelasticity, self-healing and shear-thinning properties, and controllable sol-gel transitions ^[10]^, while the dopamine groups on gelatin promote hemostasis and adhesion ^[11]^. We characterized these features of the resultant HA-PBA/Gel-Dopa hydrogel, and applied it to an animal model of brain lesions to assess its efficacy healing, including supporting cell infiltration, reducing astrogliosis, and promoting neural regeneration.

## 2. Results

### 2.1. Preparation and characterization of the HA-PBA/Gel-Dopa hydrogel

To synthesize the HA-PBA/Gel-Dopa hydrogel, the 3-aminomethyl PBA and dopamine were conjugated on HA and gelatin, respectively (**Supplementary Fig. S1**). A hydrogel with brown color was formed immediately (< 5 s) at room temperature by mixing HA-PBA and Gel-Dopa precursors in phosphate buffer solution (PBS) of pH 7.4 through the formation of boronate ester dynamic bonds between PBA and Dopa (**Fig. 1a,b**). A series of HA-PBA/Gel-Dopa hydrogels with fixed concentration of HA-PBA at 1.5% (wt/vol) and various concentrations of Gel-Dopa (3%, 4% to 5% (wt/vol)) were prepared for further characterization, which were denoted as 3% Gel-Dopa, 4% Gel-Dopa and 5% Gel-Dopa respectively.

**Figure 1.**
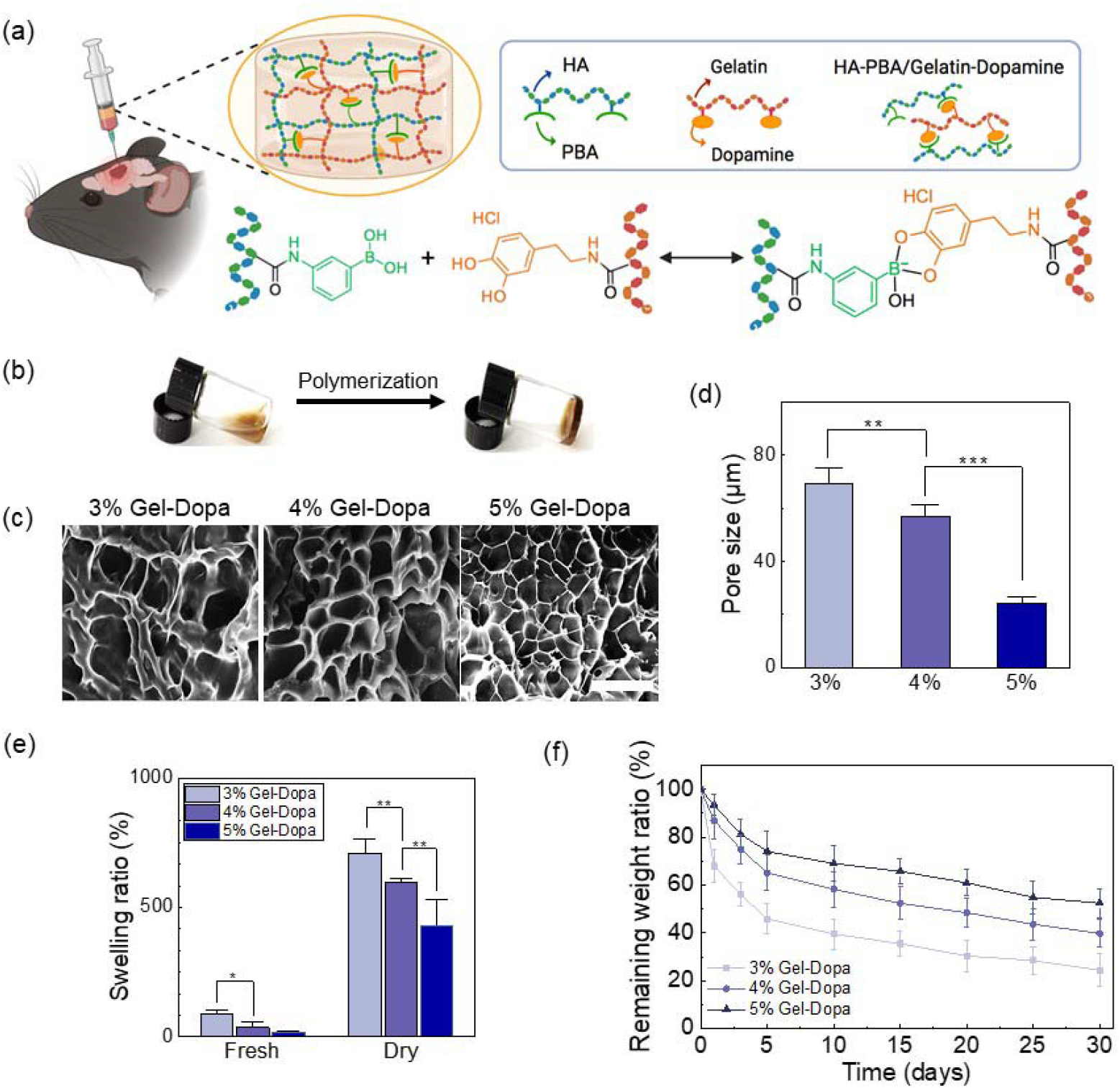
Characterization of HA-PBA/Gel-Dopa hydrogels. (a) Schematic of HA-PBA/Gel-Dopa hydrogel synthesis. (b) Gelation of HA-PBA/Gel-Dopa hydrogel. (c) SEM images of HA-PBA/Gel-Dopa hydrogel porosity (scale bar: 20 μm). (d) Pore diameters of different hydrogels after swelling. (e) Swelling ratios of fresh and air-dried hydrogels. (f) Degradation of hydrogels in PBS. * *p*<0.05, ** *p*<0.01, and *** *p*<0.001.

SEM images of HA-PBA/Gel-Dopa hydrogels showed interconnected porous structures of microscale pore size, with pore density and diameter decreasing with increasing Gel-Dopa concentration (**Fig. 1c,d**). Since swelling reflects the ability of hydrogels to absorb wound exudate, we then tested the swelling ratios of both fresh and frozen-dried hydrogels and found that all HA-PBA/Gel-Dopa hydrogels exhibited good water absorption capacity after 24 hours. Swelling ratios of frozen-dried hydrogels gradually decreased with the increase of Gel-Dopa concentration, from 86.4±17.2%, 35.6 ±20.1% to 19.3±6.4% (fresh prepared) and from 708±56.2%, 599±11.8% to 428±98.5% (frozen dried) (**Fig. 1e**). Swelling of the fresh prepared hydrogel plateaued after 24 hours (**Supplementary Fig. S2)**, a feature that is desirable for filling of a brain cavity.

The degradation rate of the HA-PBA/Gel-Dopa hydrogels could be adjusted by tuning the concentration of Gel-Dopa, which decreased with increasing Gel-Dopa concentration (**Fig. 1f**). The hydrogel with 3% Gel-Dopa showed the fastest degradation rate, remaining 26.3±6.8% of initial mass after 30 days of degradation in pH 7.4 PBS at 37 °C.

### 2.2. Rheological properties of the HA-PBA/Gel-Dopa hydrogel

We aimed to match the viscoelasticity of HA-PBA/Gel-Dopa hydrogels to that of native brain parenchyma. Rheological frequency sweep (0.1-10 rad/s) tests showed that the storage modulus (*G’*) of HA-PBA/Gel-Dopa hydrogels ranged from 30.3±10.5 Pa (3% Gel-Dopa) to 157 ± 74.5 Pa (4% Gel-Dopa) and 589±98.2 Pa (5% Gel-Dopa) (**Fig. 2a, Supplementary Fig. S3a**), which was on the order of *G’* that we measured for rat and mouse cortex (603±82.4 and 715±96.3 Pa, respectively, **Fig. 2a, Supplementary Fig. S3c,d**). The loss tangent (*G”/G’*) ranged from 0.33±0.08 (5% Gel-Dopa) to 0.46±0.15 (4% Gel-Dopa) and 0.71±0.07 (3% Gel-Dopa), also within the ranges measured for rat and mouse cortex (0.31±0.15 and 0.25±0.11, respectively, **Fig. 2b, Supplementary Fig. S3c,d**). Based on these results, we chose HA-PBA/Gel-Dopa hydrogels with 5% Gel-Dopa for all subsequent experiments.

**Figure 2.**
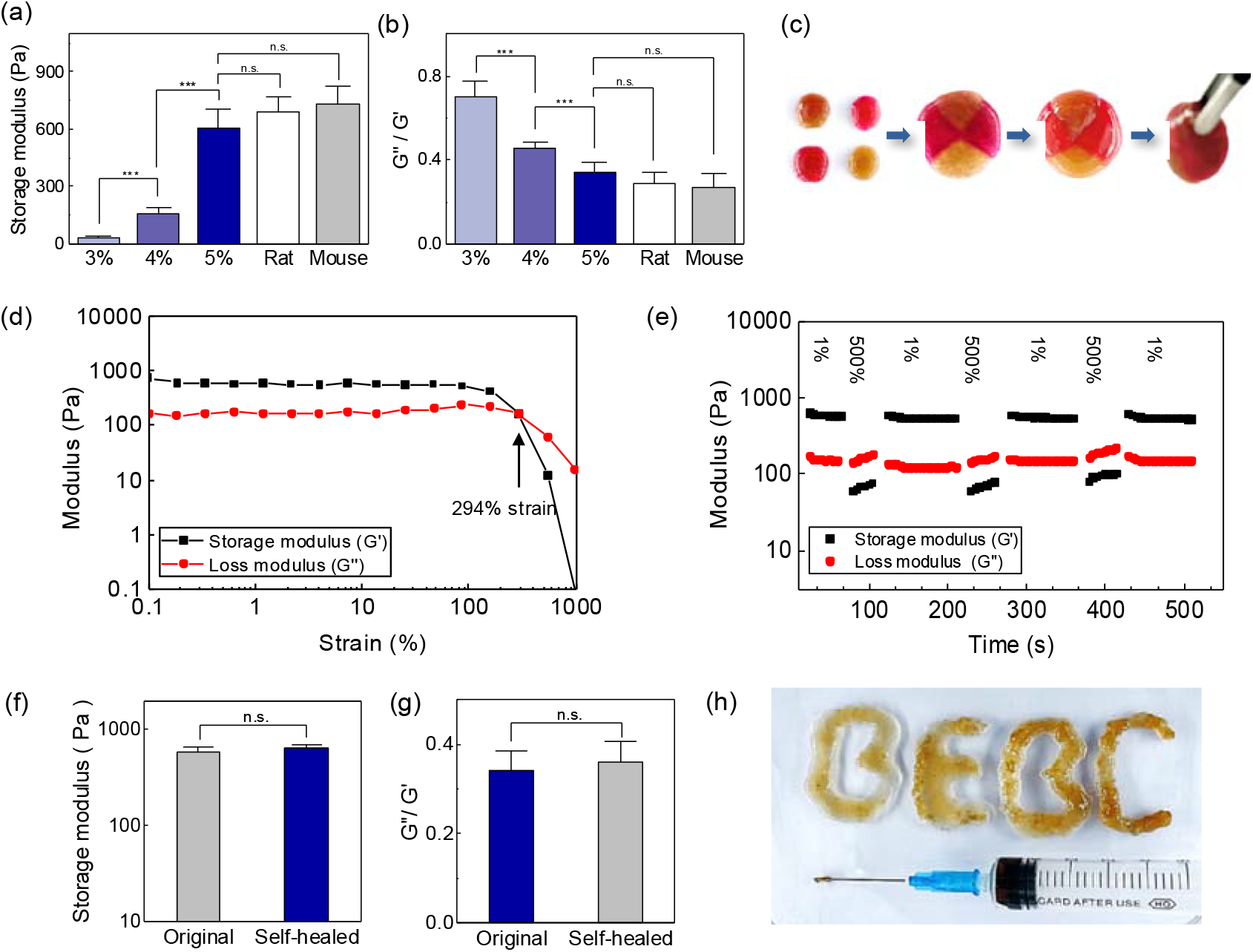
Mechanical properties of HA-PBA/Gel-Dopa hydrogels. (a) Storage modulus of different hydrogels and of rat and mouse cerebral cortex. (b) Loss tangent *G”/G’* of different hydrogels and of rat and mouse cerebral cortex. (c) Qualitative demonstration of self-healing. Shear rheometery strain amplitude sweeps of 5% Gel-Dopa hydrogel loaded at a strain rate of 1%. (e) Oscillatory loading of the 5% Gel-Dopa hydrogel at 1 Hz with a peak strain of 500% for 2 mins, followed by 1 Hz oscillations with a peak strain of 1% for 3 mins. (f, g) Storage modulus and loss tangent *G”/G’* of the original and self-healed hydrogel. (h) Qualitative demonstration of injectability of 5% Gel-Dopa hydrogel. *** *p*<0.001, n.s.: not statistically significant.

### 2.3. Self-healing and shear-thinning of the HA-PBA/Gel-Dopa hydrogel

The HA-PBA/Gel-Dopa hydrogel exhibited self-healing sufficiently strong to enable healed surfaces to be picked up with tweezers (**Fig. 2c**). To assess the self-healing capability, a series of rheological tests were performed. Strain-sweep tests showed the intersection point of *G’* and *G”* to occur at a strain of 294%, beyond which the hydrogel network would be disrupted (**Fig. 2d**). To quantify the self-healing capability, we loaded hydrogels alternately with shear strains of 500% (at which extensive disruption is expected) and of 1% (at which no disruption is expected) at a frequency of 1 Hz. *G’* decreased and *G”* increased when strain increased from 1% to 500% (**Fig. 2e**), consistent with breakdown of the hydrogel network. *G’* recovered its initial value and returned to being higher than *G”* when the strain decreased back to 1%. This value of *G’* was recovered across cyclic loadings, as was the loss tangent *G”/G’* (**Fig. 2f,g, Supplementary Fig. S3b**), confirming the self-healing capability. Shear-thinning was demonstrated by the successful injection of the HA-PBA/Gel-Dopa hydrogel through a 22 G needle, followed by quick self-healing without external intervention (**Fig. 2h**).

### 2.3. Tissue adhesive and *in vivo* hemostatic abilities of the HA-PBA/Gel-Dopa hydrogel

Adhesion, desirable to achieve wound closure and control bleeding and infection ^[12]^, was demonstrated via skin adhesion (**Fig. 3a**) and lap shear (**Fig. 3b**) tests, which demonstrated an adhesive strength of 5.38±1.12 kPa (**Fig. 3c**). To assess adhesion to brain parenchyma, we injected the hydrogel into an artificial cavity in the right cortex of a rat. The hydrogel adapted to the shape of cavity and adhered to the tissue (**Fig. 3d,e**), possibly due to hydrogen bonding at the interface ^[13]^.

**Figure 3.**
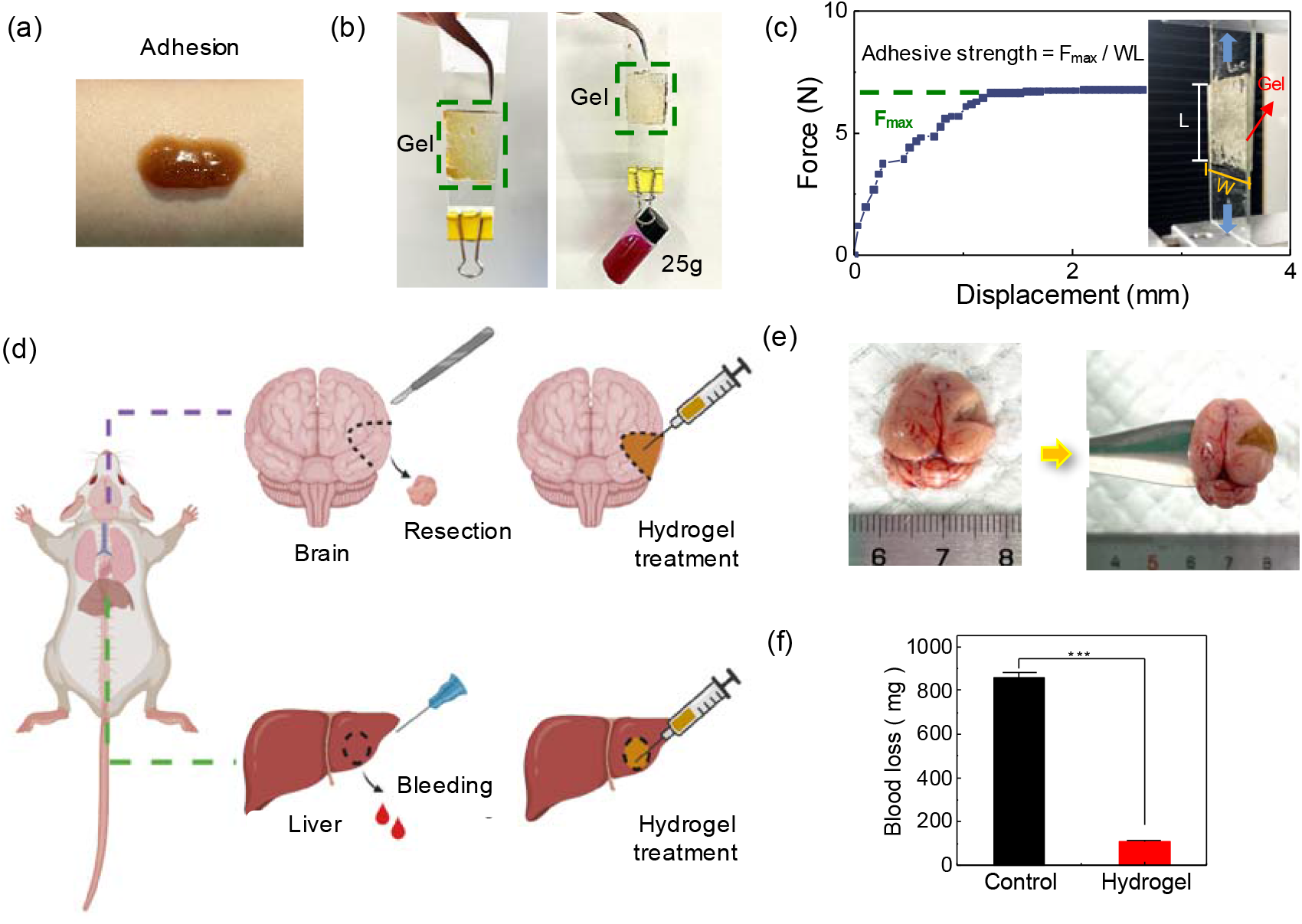
Adhesive and hemostatic properties of the HA-PBA/Gel-Dopa hydrogel. (a,b) Qualitative demonstration of adhesion to (a) skin and (b) glass. (c) Quantitative lap-shear adhesion test of 5% Gel-Dopa hydrogel. (d) Schematic representations of adhesion and hemostasis demonstrations. (e) Adhesion of 5% Gel-Dopa hydrogel to brain parenchyma. (f) The 5% Gel-Dopa hydrogel dramatically reduced blood loss. *** *p*<0.001.

Homeostasis, critical in neurosurgery and in TBI ^[14]^, was assessed in a rat liver injury model. Bleeding arising from impaling a rat liver with a 20 G needle was stopped after injection of the HA-PBA/Gel-Dopa hydrogel (**Fig. 3d,f**). Whereas 862±24.3 mg blood escaped the livers of rats in the untreated group, 107±11.5 mg escaped the livers of rats treated with the hydrogel (*p* < 0.001).

### 2.4. Biocompatibility of the HA-PBA/Gel-Dopa hydrogel

To assess the biocompatibility of the HA-PBA/Gel-Dopa hydrogel, we evaluated both the viability and spreading of primary astrocytes embedded in the hydrogel over 7 days (**Fig. 4a,b**). Viability remained high after 7 days (>98%), indicating excellent biocompatibility. The spreading area increased and the cell shape index decreased over time as reflected by F-actin staining (**Fig. 4c**), suggesting that the HA-PBA/Gel-Dopa hydrogel supported the growth of astrocytes.

**Figure 4.**
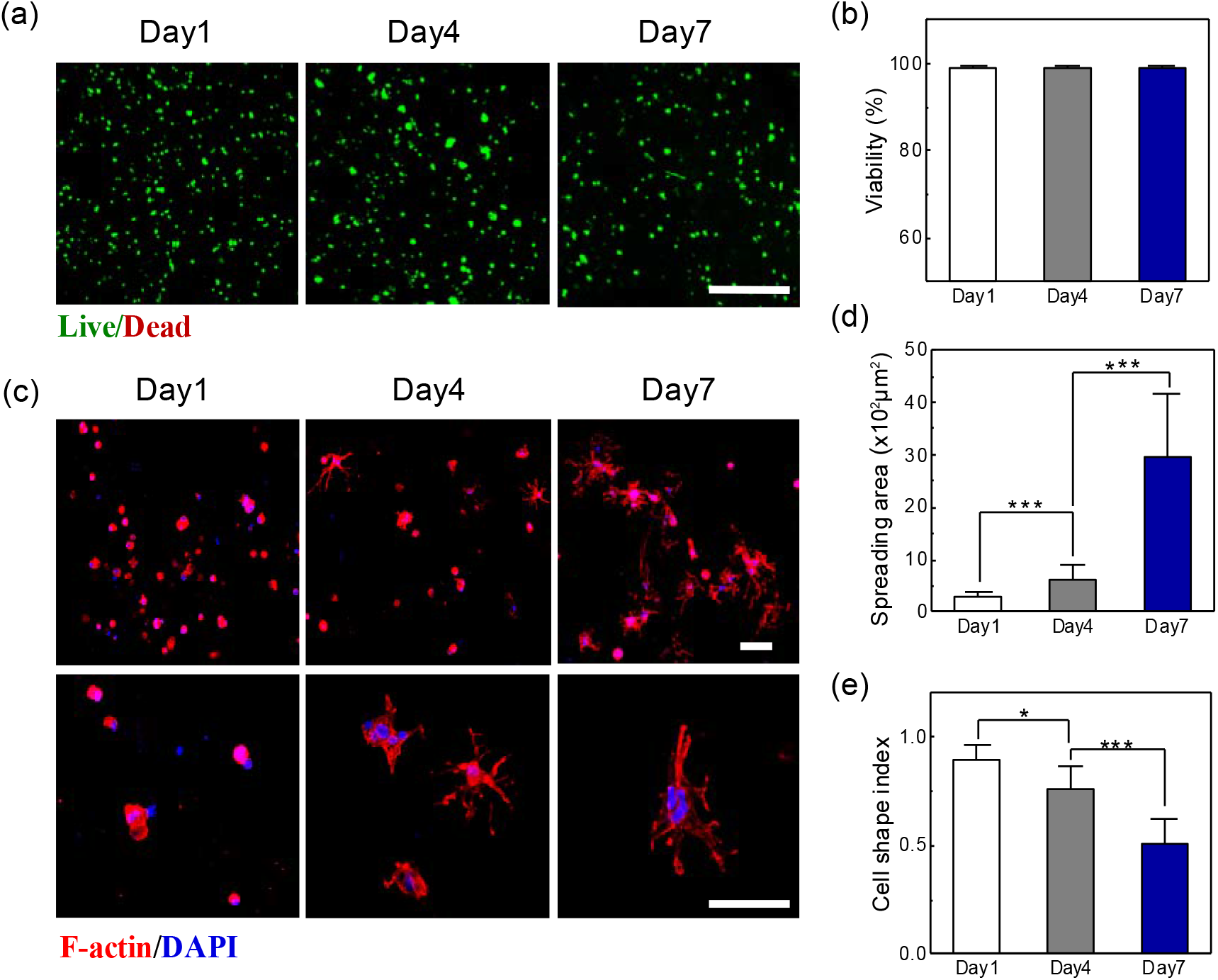
Biocompatibility of the HA-PBA/Gel-Dopa hydrogel. (a) Representative live-dead images of astrocytes cultured within 5% Gel-Dopa hydrogels after 7 days of culture. Scale bar, 200 μm. (b) Quantification of the viability of encapsulated cells calculated as the percentage of live astrocytes among all stained cells. (c) Immunofluorescence staining of encapsulated astrocytes in hydrogels after 1, 4 and 7 days of culture (red, F-actin; blue, nucleus). All scale bars, 40 μm. (d) Quantification of the spreading area of encapsulated astrocytes. (e) Quantification of the shape index of encapsulated astrocytes. * *p*<0.05 and *** *p*<0.001.

### 2.5. Histological analysis of treated brain lesions

We assessed the HA-PBA/Gel-Dopa hydrogel in a mouse brain lesion model in which a 2×2×2 mm^3^ region of the right frontal cortex was resected and then immediately filled with the HA-PBA/Gel-Dopa hydrogel (**Fig. 5a,b**). Brains of the sham injury (NS control) and hydrogel-treated groups were fixed via transcardial perfusion 7, 14 and 21 days after injury (**Fig. 5c**). Mice in the hydrogel-treated groups showed significantly less injury to the surrounding tissue than did mice in the NS control group, with hematoxylin-eosin (H&E) staining (**Fig. 5d**) and Nissl staining (**Fig. 5e**) showing substantial loss of ECM and nerve cells in the NS control group. In the NS control group, the lesion cavity expanded through the cortical layers to the white matter and the hippocampus in the ipsilateral hemisphere. In the hydrogel-treated group, the hydrogel-filled lesion generally healed and contained obvious neural cell infiltration in both the hydrogel and the lesion core. The hydrogel-treated groups showed significantly decreased lesion area and increased wound closure relative to the NS control group (**Fig. 5f,g**). Lesions nearly disappeared in the hydrogel-treated group, with a closure rate of 95.6±1.75%, persisted even at day 21 in the NS control group.

**Figure 5.**
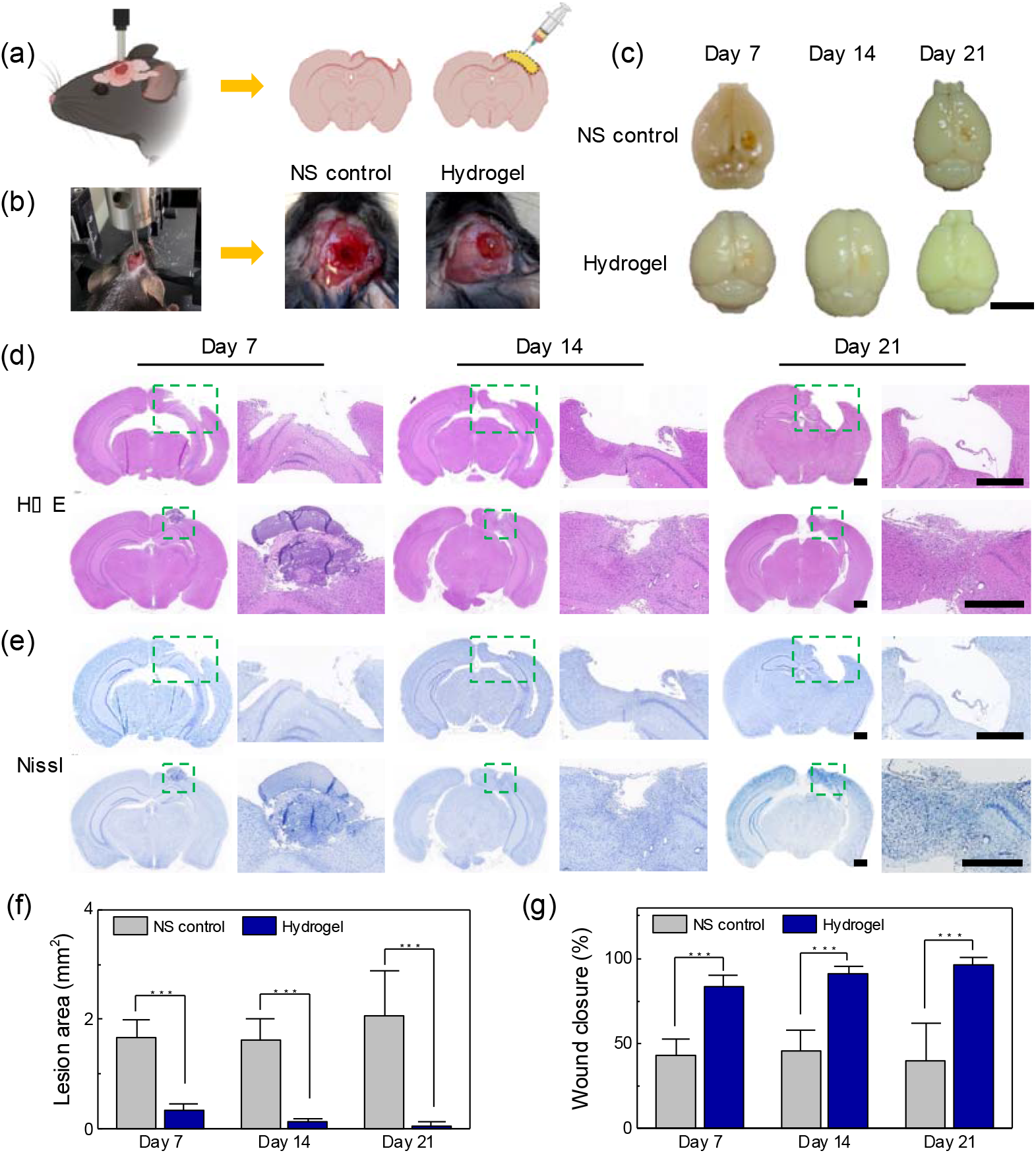
Histological analysis of regenerative brain tissue. (a) Schematics of the mouse brain lesion model. (b) Mouse brain lesion model. (c) 4% Paraformaldehyde-fixed control sham injury (NS control) and 5% Gel-Dopa hydrogel treated brains at 7, 14 and 21 days after injury. Scale bar, 5 mm. (d) H&E stains of NS control and hydrogel treated brains at 7, 14 and 21 days after injury. All scale bars, 2 mm. (e) Nissl stains of NS control and hydrogel treated brains at 7, 14 and 21 days after injury. All scale bars, 2 mm. (f) Quantification of lesion areas in the NS control and hydrogel-treatment groups at 7, 14 and 21 days after injury. Wound closure percentage in the NS control and hydrogel-treatment groups at 7, 14 and 21 days after injury. *** *p*<0.001.

We next assessed whether this support of cell infiltration and lesion cavity closure was associated with regeneration of potentially functional neural tissue. Microtomed slices of the brains of NS control and hydrogel-treated mice were fixed and stained for both microtubule-associated protein-2 (MAP-2, a specific marker of neurons) and glial fibrillary acidic protein (GFAP, a specific marker of astrocytes) at 7, 14 and 21 days after injury. Ingrowth of cells into the lesion cavity was evident at all time points in the hydrogel-treated group, but not in the NS control group (**Fig. 6**). Significant astrogliosis associated with GFAP expression was evident for all 21 days in the NS control group, but this decreased over time as the lesion closed in the hydrogel-treated group (**Fig. 6**).

**Figure 6.**
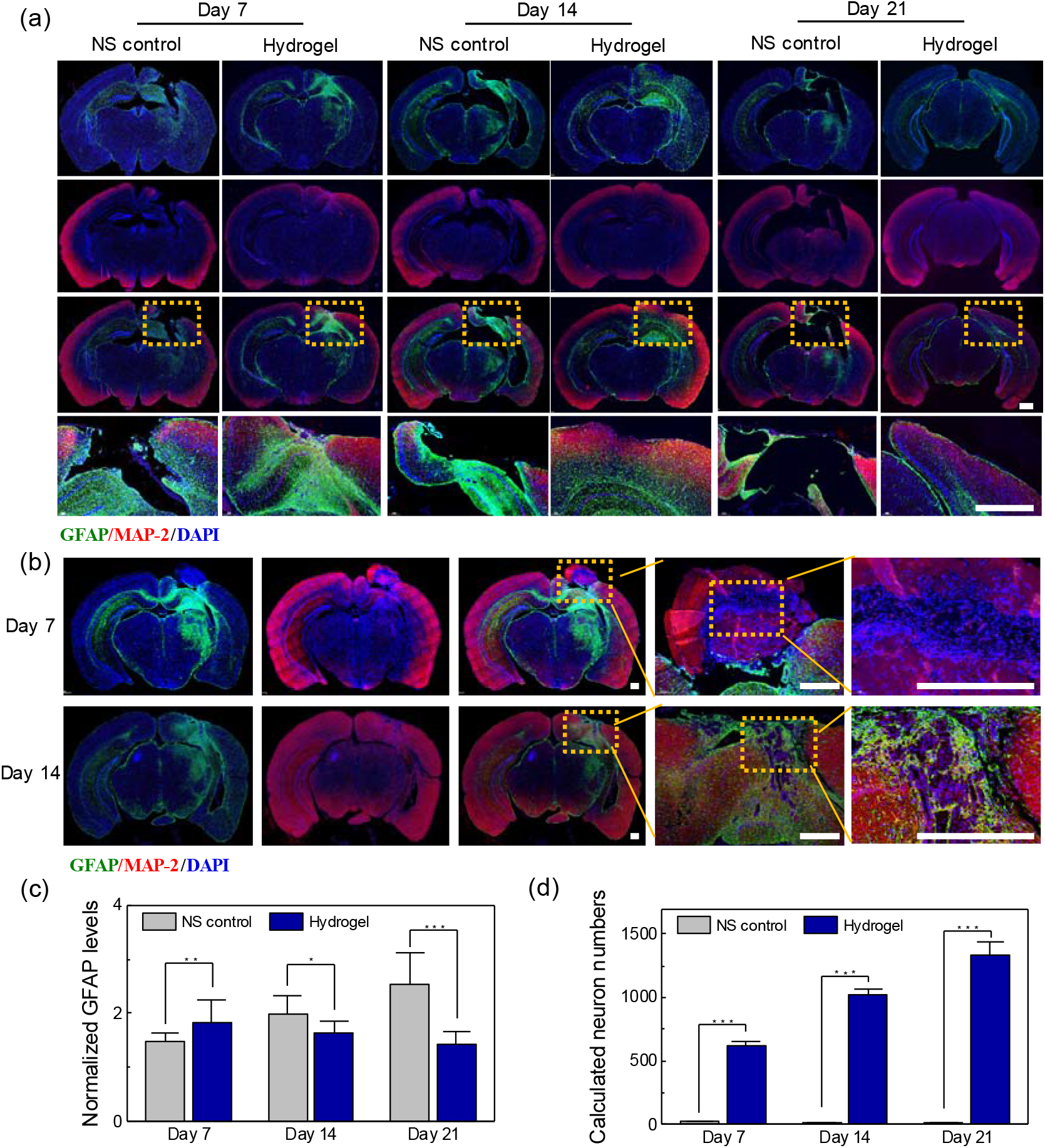
Astrogliosis surrounding the lesion cavity, and migration of neuronal cells into the lesion cavity. (a) Immunofluorescence staining of mice brain slices at 7, 14 and 21 days after injury (green, GFAP, a reporter for astrogliosis; red, MAP-2, a reporter of neurons; blue, DAPI, a stain of nuclei). All scale bars, 1mm. (b) Immunofluorescence staining of mouse brains with hydrogel-filled lesions cavity at 7 and 14 days after injury (green, GFAP; red, MAP-2; blue, DAPI). All scale bars, 500 μm. (c) Normalized GFAP levels in mouse brain slices at 7, 14 and 21 days after injury. (d) Calculated numbers of neurons in lesion cavities at 7, 14 and 21 days after injury. * p<0.05, ** p<0.01, and *** p<0.001.

## 3. Discussion

The HA-PBA/Gel-Dopa hydrogel reduced glial scar formation and enabled the ingrowth of neurons in a mouse model of a severe brain lesion. This is consistent with other reports of HA reducing glial scar formation ^[15]^. Glycosaminoglycans are the major structural and bioactive components of human brain ECM ^[5b]^, which are biocompatible, hydrophilic, largely non-immunogenic, and capable of scavenging reactive oxygen species ^[16]^. Gelatin, as denatured collagen, also mimics the components of native brain ECM ^[17]^ and is suitable for tissue engineering and regenerative medicine due to its biocompatibility, biodegradability, promotion of blood coagulation, and promotion of cell adhesion and migration ^[18]^. The introduction of catechol groups (dopamine, similar to mussel adhesion protein ^[19]^) in a hydrogel network can enhance the hydrogel’s hemostatic, antioxidant and adhesive properties ^[11]^, and may have contributed to the HA-PBA/Gel-Dopa hydrogel’s hemostatic capacity (**Fig. 3f)**. Catechol groups, which can participate in the formation of borate bonds, may also have contributed to the biodegradability of the HA-PBA/Gel-Dopa hydrogel (**Fig. 1g)**, to the viability and growth of neural cells (**Fig. 4)**, and to neural cell infiltration (**Fig. 5d,e)**.

The mechanical properties of the HA-PBA/Gel-Dopa hydrogel may also have contributed to its efficacy. The brain’s soft, viscoelastic biomechanical characteristics, which arise from non-fibrous ECM dominated by non-covalently cross-linked, highly hydrophilic, glycosaminoglycans ^[20]^, may be important to neural healing. ECM mechanics affects brain development and pathology ^[20b, 21]^ and regulates multiple cell behaviors including the astrocytic phenotype transformation ^[22]^. Soft ECM (∼40 Pa) induces astrogliosis, while stiff ECM (∼1 kPa) reverses this in a 3D *in vitro* model ^[23]^, suggesting a key role of hydrogel stiffness on glial scar progression, and underscoring our decision to match *G’* of the HA-PBA/Gel-Dopa hydrogel to that of native cortex tissues (**Fig. 2a, Supplementary Fig. S3a**).

ECM viscoelasticity may modulate nerve cell behavior and tissue homeostasis ^[7b, 24]^. This is consistent with the observation that relaxation timescales of brain tissue (∼1 min ^[25]^) match those of cellular dynamics (*e*.*g*., pseudopodia and axonal expansion timescales from tens of seconds to minutes ^[26]^). This may be important to functional cellular infiltration in neural healing. Unlike previous hydrogels for neural healing ^[6b, 27]^, the HA-PBA/Gel-Dopa hydrogel could be tuned to match the viscoelastic properties of brain parenchyma (**Fig. 2a, Supplementary Fig. S3a**), possibly affecting its therapeutic efficacy.

From the perspective of TBI, loss of ECM and neural cells from lesions can lead to a cavity that hinders cell infiltration, axonal extension and generation, and that is an obstacle to TBI recovery ^[2, 4]^. Craniotomy and infection can similarly leave patients with intracranital cavities. Because spontaneous neural regeneration does not occur ^[28]^, injectable hydrogels have been explored that have improved filling of a lesion over a 4-12 week time course ^[1a, 6b, 27]^, often with the hydrogel simultaneously delivering encapsulated stem cells or growth factors ^[6c, 29]^. Filling an intracranial cavity with our ECM-mimicking HA-PBA/Gel-Dopa hydrogel caused strong astrogliosis over the first 7 days, indicating the formation of glial scar, which resolved over the next two weeks as mature neurons entered the cavity with a structure representative of regenerated tissue (**Fig. 6a,c,d**).

Glial scars protect injured neural tissue at early stages post injury, but are in general a barrier to central nervous system regeneration in the later stages ^[30]^. However, the glial scar formed in the early stages following injection of our hydrogel did not prevent the migration of neurons into the lesion. One possible reason is that the HA-PBA/Gel-Dopa hydrogel might have induced astrocytic activation in the early stages, and another reason is that the matched viscoelastic material properties of the hydrogel might have facilitated reduction of the glial scar and invasion by neurons.

Although literature can be found that packages advances like those presented here as curing TBI or brain lesions, the field in fact has a long way to go, and we conclude by emphasizing several key caveats. First, although the tissue that appeared in the filled lesions at 21 days appeared to have the structure of cerebral cortex, further studies, including behavioral studies, need to be conducted to determine if the filled lesion is in fact functional. Second, we cannot determine whether the anatomically appropriate collections of neurons that we imaged arose from cell migration, from cell process extension, or from generation of new cells. All are possible.

Third, these experiments were performed in a mouse brain, approximately 1/60,000^th^ the volume of a human brain, and with length scales that are about 1/40^th^ that of a human. Although certain effects that we observed can be expected to scale to humans, others might not. Effects associated with cell migration and cell growth can be expected to occur over times that scale linearly with size and might thus take 40 times longer. Processes associated with diffusion occur with times that scale with the square of size, and can thus be expected to take over a thousand times longer. This would include diffusion of soluble factors to the center of the lesion in the absence of a vascular system. Processes associated with the surface chemistry of the hydrogel, such as the hemostatic aspects and possibly aspects of glial scar reduction, would not be affected by lesion size. The matching of viscoelasticity between the hydrogel and the brain parenchyma would also be independent of size, although future work must search for potential size-dependent poroelastic effects in the mechanical response of our hydrogel. For lesion closure, a range of factors that relate to volume, surface area, and length factor in, and it is impossible to determine without further experimentation how this would scale to humans, including how associated reductions in lesion volume would synergistically improve diffusion, migration, and growth-related aspects of healing. However, our results appear promising, and motivate studies in larger animals.

## 4. Conclusion

We developed an ECM-mimicking, self-healing HA-PBA/Gel-Dopa hydrogel based on a dynamic boronate ester and showed its efficacy in healing of brain lesions in a small aniomal model. The stiffness and viscoelasticity of this hydrogel could be tuned to match the mechanical properties of brain parenchyma, and the hydrogel showed effective swelling, degradation, self-healing and shear-thinning properties. Due to the dopamine groups in the network, the HA-PBA/Gel-Dopa hydrogel was adhesive and aided with hemostasis. The hydrogel supported the viability and growth of neural cells, enabled closure of lesions, and decreased glial scar formation. This study suggests a role for viscoelastic properties of hydrogels in brain lesion treatmant, including a role for a tuned cellular mechanical microenvironment, and motivates additional experimentation.

## Materials and Methods

### Synthesis of hyaluronic acid-phenylboronic acid (HA-PBA)

To synthesize HA-PBA, we used 4-(4,6-dimethoxy-1,3,5-triazin-2-yl)-4-methyl-morpholinium chloride (DMTMM, Aladdin, China) as coupling agent to conjugate 3-aminomethyl PBA (Aladdin, China) to HA (90 kDa, Melonepharma, China). In a typical reaction, 0.25 mmol HA was completely dissolved in 20 mL deionized water (DI water). Then, 0.25 mmol PBA and 0.5 mmol DMTMM was added to the HA solution separately. After all the agents were dissolved, we used 1M HCl and 1M NAOH solution to adjust the pH of the solution to 6.5. All reaction mixtures were stirred at room temperature for 24 hours. After that, the mixtures were transferred to 8 kDa molecular weight cut-off (MWCO) dialysis bags (MD, China) and dialyzed against DI water for at least 5 days at room temperature, with the water changed at least twice every day. The dialyzed solutions were freeze-dried to make HA-PBA. HA-PBA was stored at room temperature before use.

### Synthesis of gelatin-dopamine (Gel-Dopa)

To synthesize Gel-Dopa, we used ethyl-dimethyl-aminopropylcarbodiimide (EDC, Aladdin, China) and N–hydroxy-succinimide (NHS, Aladdin, China) as coupling agent to agent dopamine hydrochloride (Mackin, China) to gelatin (Type A from porcine skin, Sigma–Aldrich, USA). In a typical reaction, 1g gelatin was dissolved in 40 mL of DI water at 50 °C. EDC (1.35 g) and NHS (1.2 g) was added to the solution. After 5-10 minutes stirring, we added 1g dopamine hydrochloride. After all the agents were dissolved, we used 1M HCl and 1M NAOH solution to adjust the pH of the solution to 4.7. All reaction mixtures were stirred at 50 °C for 24 hours. Subsequently, the Gel-Dopa solution was dialyzed in DI water for 5-7 days at 50 °C and then freeze-dried.

### Chemical characterizations of HA-PBA/Gel-Dopa

1 H NMR spectra of HA-PBA/Gel-Dopa was performed by a Bruker Ascend 400MHz NMR instrument, and the solvent was deuter oxide (D2O). The HA, PBA, HA-PBA, gelatin, dopamine and Gel-Dopa were dissolved in D2O at 2 mg/mL for NMR acquisition, and the chemical shifts were referred to the solvent peak of D2O (4.78 ppm at 25 °C).

### Structural characterizations of HA-PBA/Gel-Dopa

the samples were dried by a freeze dryer, we sprayed them with a thin gold layer and examined them by a field emission scanning electron microscope (FE-SEM; QUTAN FEG 250, FEI). The pore diameters of hydrogels were measured by Image J software. For every hydrogel sample (at least 3 samples), each sample had 3-5 pictures from various regions.

### Swelling performance

Fresh group, we weighed fresh samples (*W*_*d*_) by an electronic precision balance, put them in PBS for 24 hours at room temperature, and weighed the wet weight (*W*_w_) of the samples after removing the surface moisture by filter papers. Dry group, we freeze-dried the fresh samples, weighed the dry samples (*W*_d_), put them in PBS for 24h at room temperature. Then, we weighed the wet weight (*W*_w_) of the samples. The swelling ratio of the fresh and dry samples was calculated as (*W*_w_ - *W*_d_) / *W*_d_ × 100%

### Degradation test

We prepared the samples with the same volume, removed the surface moisture of them and weighed the initial weight of them (*W*_*0*_). We Immersed the samples in PBS at constant 37 °C with shaking at 100 rpm. Then, we took out the samples, washed with DI water, dried them by an oven at 60 °C and weighed the samples at the predetermined time point (*W*_*t*_). The remaining weight of hydrogel (%) was calculated as *W*_t_ / *W*_0_ × 100%

### Mechanical characterization

Samples were prepared as discs with a diameter of 15 mm and a thickness of 1 mm. For frequency sweep test, hydrogels were tested from 0.1 rad/s to 100 rad/s with a 1% constant strain at 25□. We used 5%Gel-Dopa as an example to evaluate the self-healing property. To find the critical stain for gel failure, we conducted oscillatory strain amplitude sweep from 0.1% to 1000% at a constant frequency of 1 Hz. Then, we performed the step-stain sweep by repeating large strain (550%, 2 minutes) for network disruption and small stain (1%, 3 minutes) for mechanical recovery.

### Mechanical test for adhesives

The adhered samples were lap-shear tested Put the 5%Gel-Dopa samples between two pieces of clean and smooth glass, press by hand for 10s to adhere. For each test, the actual bonding area on the adhered substrates was individually measured. The tensile rate was 5 mm/min. Adhesive strength = *F*_*max*_ */ WL*, where F_max_ is the max value of force, W is the width of bonding area and L is the length of bonding area.

### In vivo hemostatic ability test

All animals were obtained from the Laboratory Animal Center of Xi’an Jiaotong University Health Science Center, China. All the procedures and ethics guidelines were approved by the Committee for Experimental Animal Use and Care of Xi’an Jiaotong University Health Science Center, China. We used hemorrhaging liver Sprague-Dawley (SD) rats (250 g, male) to evaluate the hemostatic ability of the 1.5% (wt/vol) HA-PBA/ 5% (wt/vol) Gel-Dopa hydrogel. We fixed the anesthetized rat on a surgical corkboard and exposed the liver of the rat by abdominal incision. We removed the liquid around the liver and placed a pre-weighted filter paper beneath the liver. Then, we stabbed the rat liver with a 20G needle, causing immediate massive bleeding. For hydrogel-treated group, we applied 150 μL hydrogel on the injury site. After 5 minutes, we weighed the filter paper with absorbed blood and compared with the control group (no treatment).

### 3D astrocyte culture

Primary astrocytes were isolated from the cortex of neonatal SD rats (1-day-old). All experiments were performed using astrocytes from the first passage (P1). P1 cells were seeded at a density of 1 × 10^6^cells/mL in 5%Gel-Dopa hydrogel and were cultured in DMEM/F12 medium containing 10 % FBS and 1% Pen-Strep for 7days.

### Mouse brain injury modeling and Histological Studies

We took C57BL/6J male mice for this study. First, we divided the mice into two groups, with 15 mice in each group. After anesthetized them by intraperitoneal injection of 2% Avertin at 10 mg/mL dosage, we shaved the hair of the mice. Then, we made a precise midline sagittal incision to expose the skull of the mice. We placed a cylindrical rod of 2 mm diameter on the exposed left parietal skull and let it drop at a speed of 2.5 m/s for 0.2 second to make a 2 mm depth injury. For the hydrogel-treated group, we immediately filled 1.5% (wt/vol) HA-PBA/ 5% (wt/vol) Gel-Dopa hydrogel (100μL) into the cavity, and closed the wounds using a suture needle. Meanwhile, for the control group (NS control), we just closed the wounds. We killed every 4 mice in each group by transcardial perfusion using 4% formaldehyde at the 7^th^, 14^th^ and 21^st^ day after injury (n = 5). After perfusion, the brains were isolated, fixed, and processed for histological studies. coronal brain sections of mice were performed with HE staining, Nissl staining and GFAP/MAP-2 double immunofluorescence staining.

### Immunofluorescence staining

We fixed the samples by 4% paraformaldehyde for 20min and washed them with PBS, penetrated the samples with 0.5% Triton-X100 (Sigma, USA) for 15min and washed them with PBS. Then we treated the samples with PBS containing 5% bovine serum albumin (BSA; MP Biomedicals, USA) and 10% normal goat serum (Thermo Fisher Scientific, USA) for 1h. The samples were treated with primary antibodies prepared in proper dilutions in PBS containing 1% BSA overnight at 4□ and were washed with PBS. We incubated the samples with appropriate Alexa 488 or Alexa 594 secondary antibodies (1:400, Thermo Fisher Scientific, USA). To visualize F-actin, samples was stained by Alexa 488 labeled phalloidin for 1 hour. The following primary antibodies were used for immunofluorescence: mouse anti-GFAP (1:400; Cell signaling Technology, 3670, USA) and rabbit anti-MAP-2 (1:300; Bioss, bs-1369R, China). After labeling nuclei with DAPI (1 μg/mL; Sigma, D9542, USA) for 10 minutes, fluorescent images were taken by an Olympus FV3000 Laser scanning confocal microscope.

### Characterization of cell viability

We incubated the samples in 1mL PBS containing 0.5 μL Calcein-AM (Life Technologies, USA) and 2 μL EthD-1 (Life Technologies, USA) at 37°C for 15 minutes, washed them with PBS, and took pictures with the Olympus FV3000 Laser scanning confocal microscope. The amount of live and dead cells was counted by a cell counter plug-in for ImageJ.

### Cell shape index

Based on the F-actin immunofluorescence staining of samples, the cell shape index was analyzed by Fuji software (ImageJ), the higher the cell shape index, the closer the cell tends to the circle. C = 4 pi A/P^2^ ∼ 12.57 A/P^2^, where C is the circularity, A is the area and P is the perimeter.

### Characterization of normalized GFAP levels

The quantitative analysis of GFAP was based on the scans. We set the fluorescence intensity of control area (left part) in each sample to the same parameter (as “1”) and compared the fluorescence intensity of right injury area with the homologous area in left control part. The normalized protein levels of samples were analyzed by Fuji software with Image plugin.

### Lesion area and wound closure

We measured the lesion area by Fuji software (ImageJ) based on the scans of HE and Nissl staining. And the wound closure (%) was calculated as *(Area(D*_*0*_*)-Area(D*_*n*_) / *Area(D*_*0*_) × 100%, *Area(D*_*0*_), where “n” represents the predetermined time point.

### Statistical analysis

Statistical analysis was conducted by using the OriginPro software package (OriginPro 2018; Origin Lab, Northampton, USA). Statistics were presented as a mean ± standard deviation for all quantitative data. One-way analysis of variance (ANOVA) was used for two-group comparison along with a t-test (\**p* < 0.05, \*\**p* < 0.01, and \*\*\**p* < 0.001).

## Acknowledgements

This work was financially supported by the National Natural Science Foundation of China (11972280, 11761161004, 12002263), the Young Talent Support Plan of Xi’an Jiaotong University, the Fundamental Research Funds for the Central Universities (xzy012020079, xzd012021037), and the Opening Project of Key Laboratory of Shaanxi Province for Craniofacial Precision Medicine Research, College of Stomatology, Xi’an Jiaotong University (2020LHM-KFKT005).

