## Supplementary material for "A Self-Healing, Viscoelastic Hydrogel Promotes Healing of Brain Lesions": supplymental

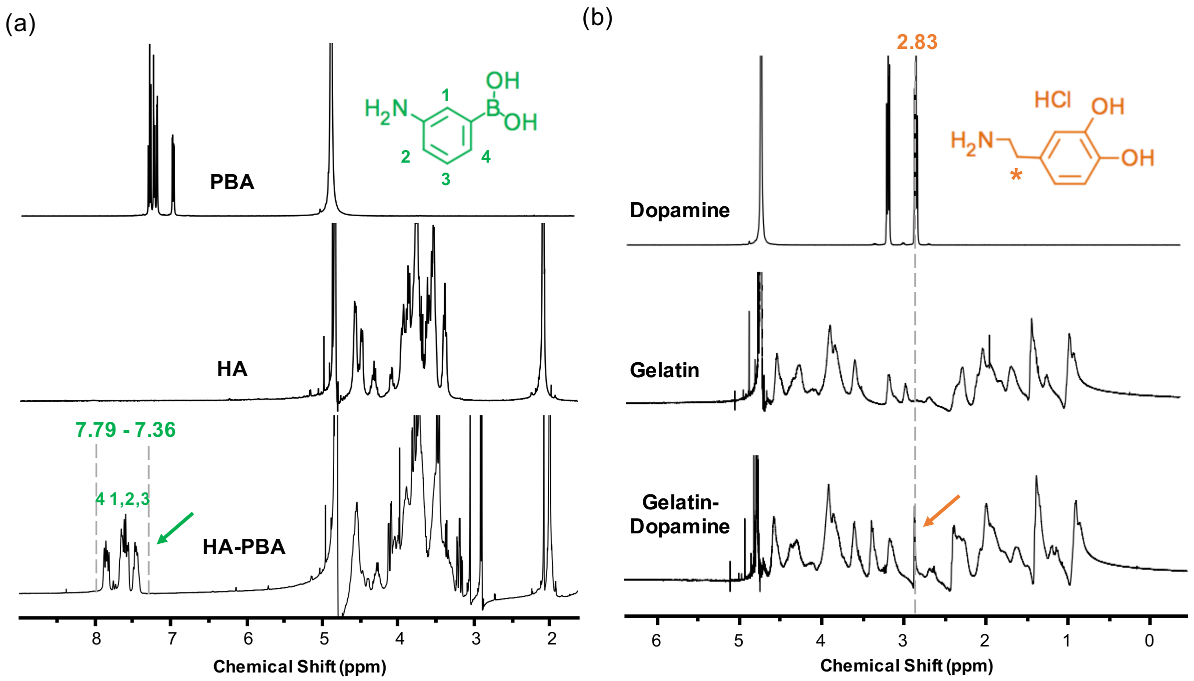


**Supplementary Figure S1.** (a) NMR spectra of hyaluronic acid (HA) and HA-PBA. (b) NMR spectra of gelatin and gelatin-Dopamine.

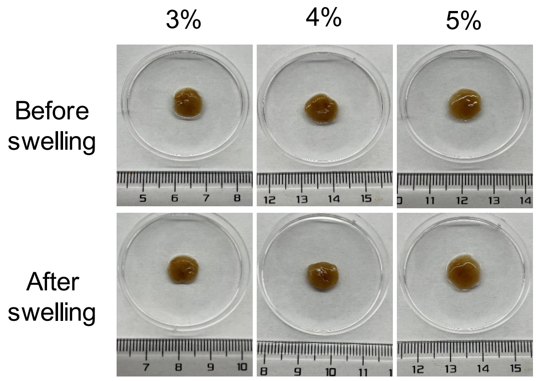


**Supplementary Figure S2.** The fresh hydrogels before and after swelling for 24 hours.

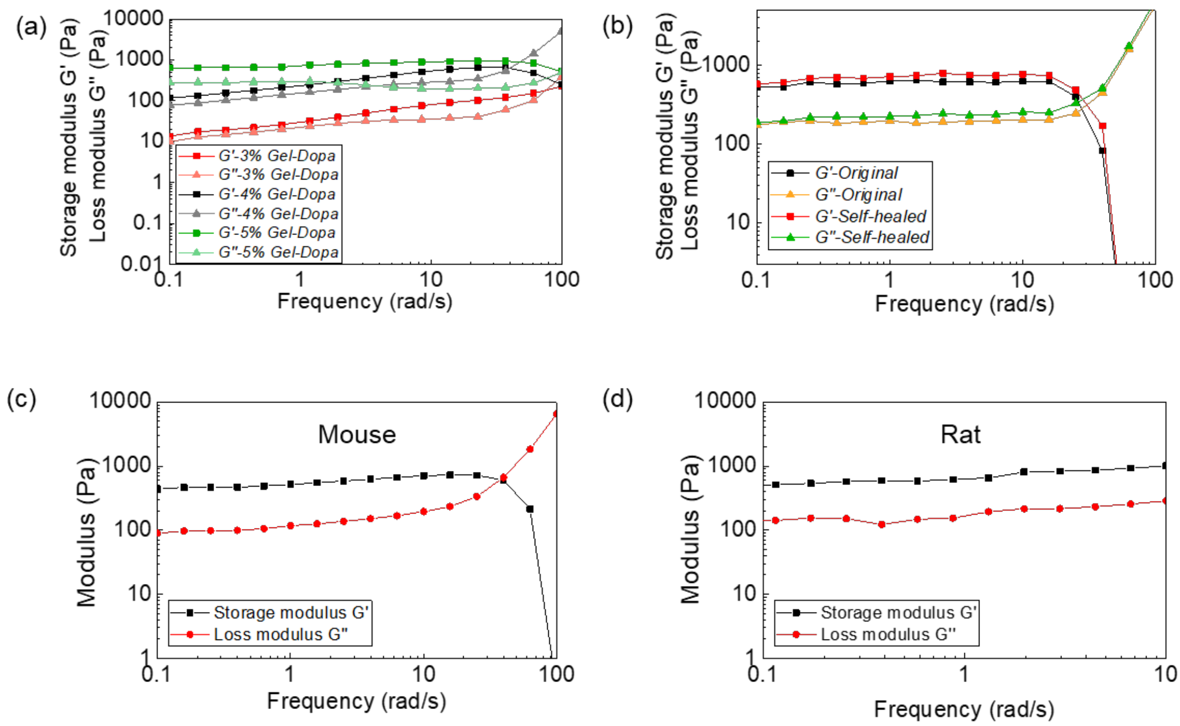


**Supplementary Figure S3.** (a) The storage modulus *G'* and loss modulus *G"* on frequency sweep of HA-PBA/Gel-Dopa hydrogels. (b) Storage modulus G' and loss modulus *G"* of original hydrogel and self-healing hydrogel on frequency sweep (c) Storage modulus *G'* and loss modulus *G"* of mouse cortex tissue on frequency sweep. (d) Storage modulus *G'* and loss modulus *G"* of rat cortex tissue on frequency sweep.
